# A prognostic signature for lower-grade gliomas based on expression of long noncoding RNAs

**DOI:** 10.1101/442616

**Authors:** Manjari Kiran, Ajay Chatrath, Xiwei Tang, Daniel Macrae Keenan, Anindya Dutta

## Abstract

Diffuse low-grade and intermediate-grade gliomas (together known as lower-grade gliomas, WHO grade II and III) develop in the supporting glial cells of brain and are the most common types of primary brain tumor. Despite a better prognosis for lower-grade gliomas, 70% of patients undergo high-grade transformation within 10 years, stressing the importance of better prognosis. Long non-coding RNAs (lncRNAs) are gaining attention as potential biomarkers for cancer diagnosis and prognosis. We have developed a computational model, UVA8, for prognosis of lower-grade gliomas by combining lncRNA expression, Cox regression and L1-LASSO penalization. The model was trained on a subset of patients in TCGA. Patients in TCGA, as well as a completely independent validation set (CGGA) could be dichotomized based on their risk score, a linear combination of the level of each prognostic lncRNA weighted by its multivariable cox regression coefficient. UVA8 is an independent predictor of survival and outperforms standard epidemiological approaches and previous published lncRNA-based predictors as a survival model. Guilt-by-association studies of the lncRNAs in UVA8, all of which predict good outcome, suggest they have a role in suppressing interferon stimulated response and epithelial to mesenchymal transition. The expression levels of 8 lncRNAs can be combined to produce a prognostic tool applicable to diverse populations of glioma patients. The 8 lncRNA (UVA8) based score can identify grade II and grade III glioma patients with poor outcome and thus identify patients who should receive more aggressive therapy at the outset.

## Introduction

Over the past decade, high-throughput RNA-seq technology discovered many novel transcriptional units, which were otherwise missed by probe design based transcriptome profiling. Among these transcriptional units were many long non-coding RNAs (lncRNA), which are transcripts longer than 200 bases with almost no protein-coding potential or open reading frames of <50 amino acids. These lncRNAs are numerous in cells [1], are highly regulated and are more cell-type specific than protein-coding genes [2]. LncRNAs are involved in a broad spectrum of function and recent studies suggest they have specific roles in different diseases like cancer (reviewed in [3, 4]).

Gliomas are the most common form of primary malignant brain tumor, which originate in the supporting glial cells in the brain, including astrocytes, oligodendrocytes and ependymal cells. Based on WHO 2016 grading system, gliomas are classified into lower-grade and much aggressive high-grade gliomas. Grade I is mostly benign, whereas diffuse low-grade and intermediate-grade gliomas make up the WHO grade II and III lesions. Grade IV gliomas include secondary glioblastomas (derived from lower grade gliomas) and primary glioblastoma multiforme (GBM). Surgical resection of tumor is the most common initial treatment for gliomas followed by radiation therapy and chemotherapy, which can increase survival to 12 months [5, 6]. Molecular markers like 1p/19q co-deletion, MGMT promoter methylation and mutation in IDH1 gene are strong predictors of survival for gliomas [7]. Lower-grade gliomas have a better prognosis than high-grade gliomas. Despite a better prognosis for lower-grade gliomas than the grade IV tumors, 70% of patients from the former group undergo high-grade transformation within 10 years.

LncRNAs are widely expressed in the central nervous system (CNS) and are involved in several pathways related to CNS development [8–13]. LncRNA BRN1B is one of the critical lncRNAs for brain development [13]. LncRNA Sox2OT plays an important role in determining neural fate [14]. Dysregulation of many lncRNAs like DGCR5, NRON, H19, DISC2 have been associated with different CNS diseases [15–18]. Previous studies have shown that specific lncRNA expression patterns are also associated with different histological subtypes and grade in gliomas [19, 20]. For example, expression of MALAT1, POU3F3 and H19 are highly correlated with glioma malignancy. More recently, lncRNAs are also found to be of prognostic significance suggesting their role in glioma malignancies and as a potential therapeutic target and biomarker [19, 20]. Li et al, 2014 revealed three molecular subtypes of gliomas based on lncRNAs expression that has a strong correlation with patient’s survival [21]. Furthermore, analysis on previously published microarray data has explored lncRNA-based signature as a prognostic marker in gliomas ([20, 22–25]).

Many studies have highlighted the power of gene expression profiles to predict tumor classification, patient outcome and tumor response to therapy. Differentially expressed genes in cancer patients versus normal individuals are often the starting set to predict prognostic signature associated with survival. This strategy suffers from false negatives and from the fact that differentially expressed genes might not be associated with differences in survival at all. Another limitation of this method is the requirement of perfect matched normal to identify differentially expressed genes. This creates a major hurdle in case of brain cancer where getting a perfect matched normal tissue is not trivial. While high-throughput technologies have facilitated the search of biomarkers through multivariate data analyses, there still remain challenges with respect to meaningful statistical and biological information. Firstly, most of the biological datasets suffer with multicollinearity: the influence of one gene on expression of other genes. Secondly, there are more features (genes) than observations (patients), which leads to overfitting by most of existing learning algorithms and results in poor performance of the model in prediction in an unseen testing dataset. Thus, a more robust machine learning approach is required to find genes as prognostic signature from a multi-dimensional multivariate gene-expression data. Regression models like lasso, ridge and elastic net are some widely used approaches to penalize the effect of multicollinearity and are well suited for constructing models when there are large numbers of features.

In the present study, we develop an lncRNA-based prognostic signature in combination with Cox regression and L1-LASSO regularization to model survival of grade II and grade III glioma patients. This is the first study that combined Cox and Lasso regularization to select lncRNAs that can predict survival in glioma patients. After controlling for covariates associated with glioma survival (age, grade, IDH1 mutation status), we selected 8 lncRNAs UVA8, to calculate a risk-score, which successfully divides patients into high-risk and low-risk groups in both TCGA (461 patients) and CGGA (274 patients) dataset. The risk score calculated by these 8 lncRNAs is an independent and better prognostic marker for grade II and grade III glioma patient survival. The guilt-by-association analysis of lncRNAs in UVA8 indicated their role in suppressing interferon signaling pathway and epithelial to mesenchymal transition. Besides their use as a biomarker, these lncRNAs need to be studied in detail to determine how they affect patient outcome.

## Materials and Methods

### Patients and samples

Aligned bam files and clinical information for 512 LGG patients (grade II and III) were retrieved from The Cancer Genome Atlas (TCGA) data portal https://portal.gdc.cancer.gov/. The study is performed on 461 patients for which both RNAseq and survival information were available. Most samples in TCGA are collected from patients from the US and also from other countries including Canada, Russia, and Italy. This dataset being the largest and most updated glioma dataset is used as training dataset in the present study. The raw sequencing data for 274 glioma patients from Chinese Glioma Genome Atlas (CGGA) as independent cohort was downloaded using accession no. SRP027383 [26]. The survival information for these Chinese patients was downloaded from CGGA http://www.cgga.org.cn/. IDH1 mutation data for all the LGG patients were retrieved from Tier 3 TCGA data accessed from the Broad GDAC Firehose; https://gdac.broadinstitute.org.

### RNASeq data quantification and analysis

The most recent version of Gencode (GENCODE v 26) GTF file available at the time of this study was used for the gene quantification [27]. Gene abundance in FPKM was obtained for 58219 genes with 15787 genes annotated as lncRNA in GENCODE v26 using Stringtie v1.3.3 [28]. Out of 15787 lncRNAs, 1289 lncRNAs with a median expression of 1 FPKM in 512 LGG patients were finally considered for the survival model.

### Survival model selection process

The gene-expression data for lncRNAs was Z-score transformed to avoid systematic error across different experiments. We first randomly selected 60% of TCGA patients for training set and remaining 40% of TCGA patients for testing set. Since, clinical information like age, gender, tumor grade or IDH mutation status can have an effect on survival (**Figure S1**), we assessed the prognostic potential of each lncRNA by multivariate Cox-regression controlling the effects from these other variables. We used FDR corrected p-value cutoff of 0.05 obtained after log-likelihood test comparing restricted (Age, Gender, tumor grade and IDH mutation status) with unrestricted (lncRNA expression, Age, Gender, tumor grade and IDH mutation status) model to identify the significant association of an lncRNA with survival. We used Cox-Proportional Hazards model based on L1 – penalized (LASSO) estimation to select the best model comprising a subset of prognostic lncRNA [29–31]. We used LASSO because it is suited for constructing models when there is a large number of correlated covariates [30].

### Risk Score calculation

Risk score for each patient was established by including each of the selected genes weighted by their estimated regression coefficients in the multivariable Cox regression analysis as discussed in previous studies [32, 33].

UVA8 Risk score = (−0.378 x expression value of RP11-266K4.14) + (−0.301 x expression value of FLJ37035)+ (−0.280 x expression value of LINC01561) + (−0.368 x expression value of RP11-118K6.3) + (−0.369 x expression value of DGCR9) + (−0.299 x expression value of RP11-142A22.3) + (−0.434 x expression value of LINC00641) + (−0.543 x expression value of RP11-96H19.1).

Coefficients are median cox-coefficient (after lasso selection and multivariate cox-regression) for each of the 8 lncRNAs from the successful models (models which can stratify patients in testing set).

### Statistical Analysis

R package glmnet was used to perform L1-penalized cox regression [34]. R package survival and survminer were used for survival data analysis and generating Kaplan–Meier plots. Different survival models were compared by time-dependent concordance index (Cindex) [35]. Cindex is the most commonly used performance measure for survival models, which calculates the fraction of pairs whose predicted survival time is correctly ordered. R package pec::cindex is used to calculate time dependent cindex [36].

## Results

### Building the lncRNA based survival model

We developed an lncRNA based survival model for gliomas through the following steps **(Figure 1).**

1) We first randomly selected 60% (n=277) of the patients from TCGA as training set and reserved the remaining 40% (n=184) of patients as testing set. The results remain similar with 70% patients in training and 30% in testing set (**Figure S3 A**).
2) Cox multivariate regression was carried out in the training set on 1289 lncRNA controlling for effects from other covariates like age, gender, tumor grade and IDH1 mutation status.
3) LncRNAs significantly associated with survival after likelihood ratio test (FDR p□<□0.05) were retained for selecting lncRNAs by lasso regularization.
4) After lasso regularization and lncRNA selection, a risk score formula was established by including selected lncRNAs weighted by their estimated regression coefficients in the multivariable Cox regression analysis. Risk Score 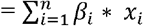 (where, □ is coefficient and *x* is expression level of lncRNA *i*)
5) Patients were classified into high-risk and low-risk group by using the median risk score as the cutoff in the training set. The coefficient for each lncRNA and cutoff of risk score obtained from training set was used to calculate risk score and stratify patients into two groups in testing set.
6) Survival differences between the low-risk and high-risk groups in the training and testing sets were assessed by the Kaplan–Meier estimate and compared using the log-rank test.

Steps 1-6 were repeated 100 times to obtain up to 100 different lncRNA subsets (models). Only those models that separated patients in the testing set such that those with low-risk score had significantly better survival than those with high-risk score were considered as successful models and retained.

**Figure 1.**
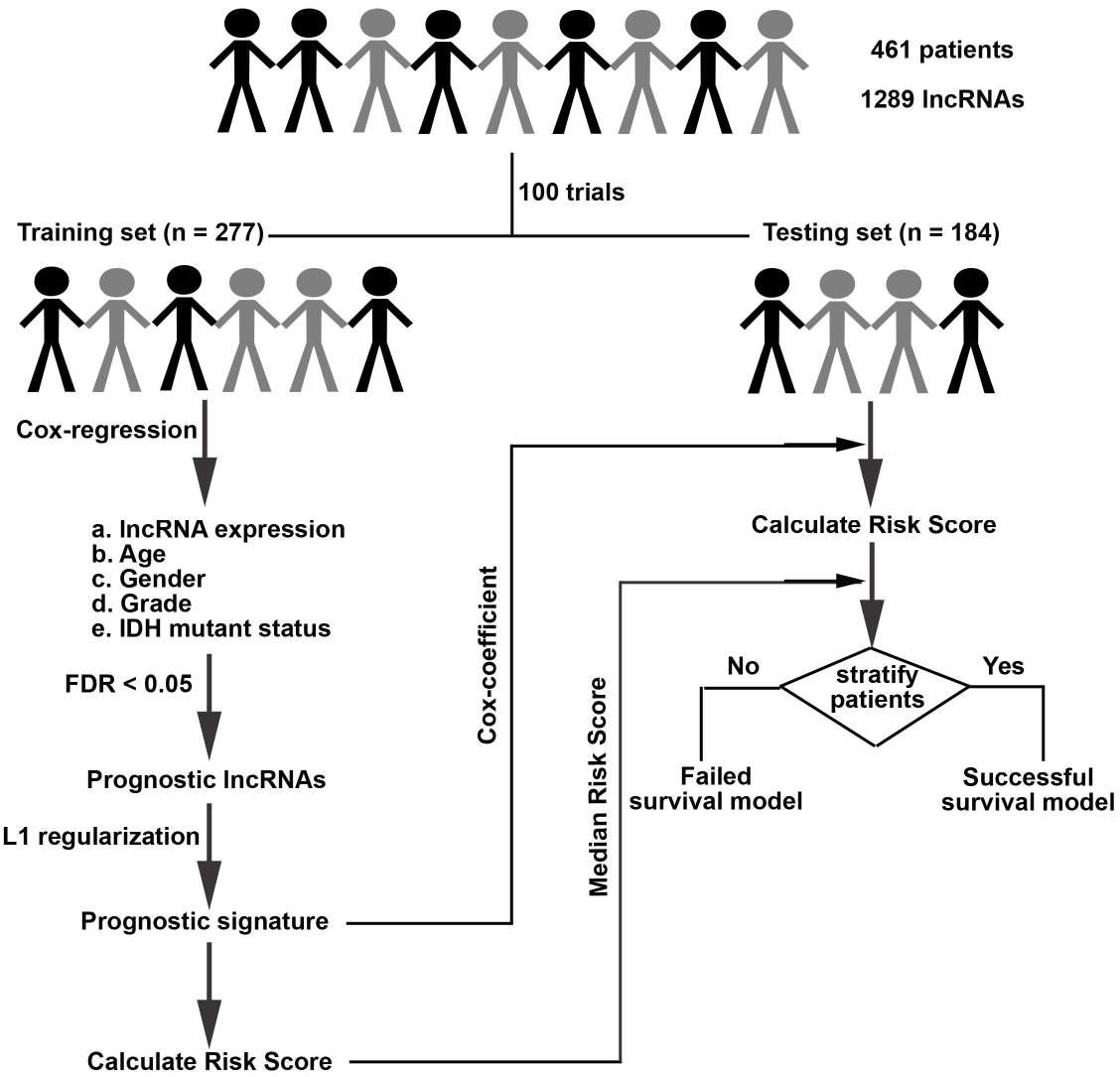
Flowchart showing steps involved in identification of lncRNA based prognostic signature.

The result obtained from one such survival model is shown in **FigureS2**. In ~20% of the trials the multivariate cox-regression and lasso regularization in the training set did not select any lncRNAs significantly associated with survival (NA in **Figure 2A**). The remaining 80% of the survival models contained different numbers of lncRNAs (x-axis of **Figure 2A**) that significantly stratify patients into low and high-risk groups in training set (**Figure 2A**). Among these 80% of survival models, 86% also significantly separated patients into high-risk and low-risk in the testing set and are referred to as successful survival models. In order to create a robust survival model we sorted the lncRNAs based on the number of times an lncRNA was selected by successful survival models (**Figure 2B**). Out of 167 total prognostic lncRNA in 69 successful survival models, we first ranked lncRNAs based on number of times a given RNA was selected by successful models and then from the top 20 selected 8 lncRNAs with the highest median cox-coefficient (Absolute value > 0.2) and least variance in the successful models in the testing set (Absolute value < 0.10). 7 out of these 8 lncRNAs were also selected after 70%-30% split of training and testing patients (**Figure S3A**), after 1000 trials instead of 100 (**Figure S3B**) and all 8 lncRNAs were selected when we used Elastic net, instead of Lasso, for regularization and lncRNA selection (**Figure S3C**) suggesting the prognostic importance of these 8 lncRNAs in gliomas. For brevity, this set of 8 lncRNA as a prognostic signature of gliomas will be referred to as UVA8 in the manuscript.

**Figure 2.**
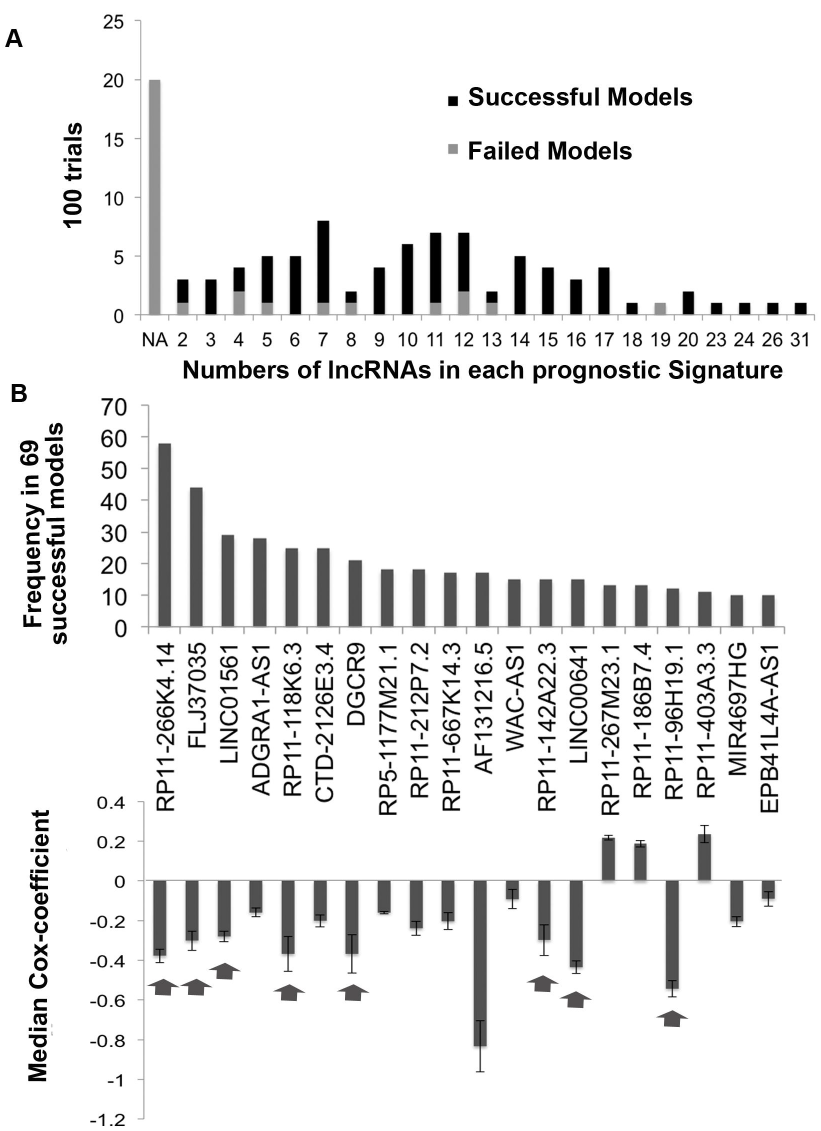
Selection of lncRNAs with best predictors of outcome. **A)** Barplot showing number of lncRNAs that predicted outcome in the training set in 100 trials. The successful models were those that also predicted outcome in the testing set. NA: no lncRNA predicted outcome in training set. **B)** Barplot showing number of times each of the top 20 lncRNAs (out of 167) were present in successful survival models (significant in testing set). The lower panel shows median Cox-coefficient (after lasso penalization) and the variance of the cox-coefficient for each of the above 20 lncRNAs from the successful models where they were selected. The arrow points towards lncRNAs selected for UVA8.

### UVA8 is predictive of survival in training and independent validation set

We assessed the predictive power of UVA8 by comparing overall survival of low and high risk patients in the entire TCGA dataset stratified based on median risk score obtained by UVA8 (risk score calculation discussed in methods). Patients in the low-risk group showed longer overall survival than the high-risk group in TCGA dataset **(Figure 3A**, median OS 741.5 vs 639 days; P = 3.1e-15, HR=5.8). The risk scores of the patients in the TCGA dataset range from −4 to 4 with median risk score of −0.023 (**Figure 3B,** top panel). Moreover, there are more patients alive in the low risk group than in the high-risk group (**Figure 3B,** middle panel). Interestingly, expression levels of all lncRNA in UVA8 are high in low risk patients than in high-risk patients indicating these lncRNAs as favorable prognostic genes ((**Figure 3B,** bottom panel)). These findings were further validated in an independent validation dataset comprising of 274 patients obtained from CGGA. Using the same median coefficient of UVA8 obtained from the successful survival models in TCGA, patients showed longer overall survival in low-risk than in high-risk group in CGGA (**Figure 3C**, median OS = 1120.5 vs 587 days; P = 0.0017, HR=1.68). Moreover, low-risk group in CGGA has also longer progression free survival (PFS) than the high-risk group (**Figure 3D,** median PFS 597.5 vs 411.5 days; P = 0.00088, HR=1.70). Thus, UVA8 can predict survival in both training and independent validation set.

**Figure 3.**
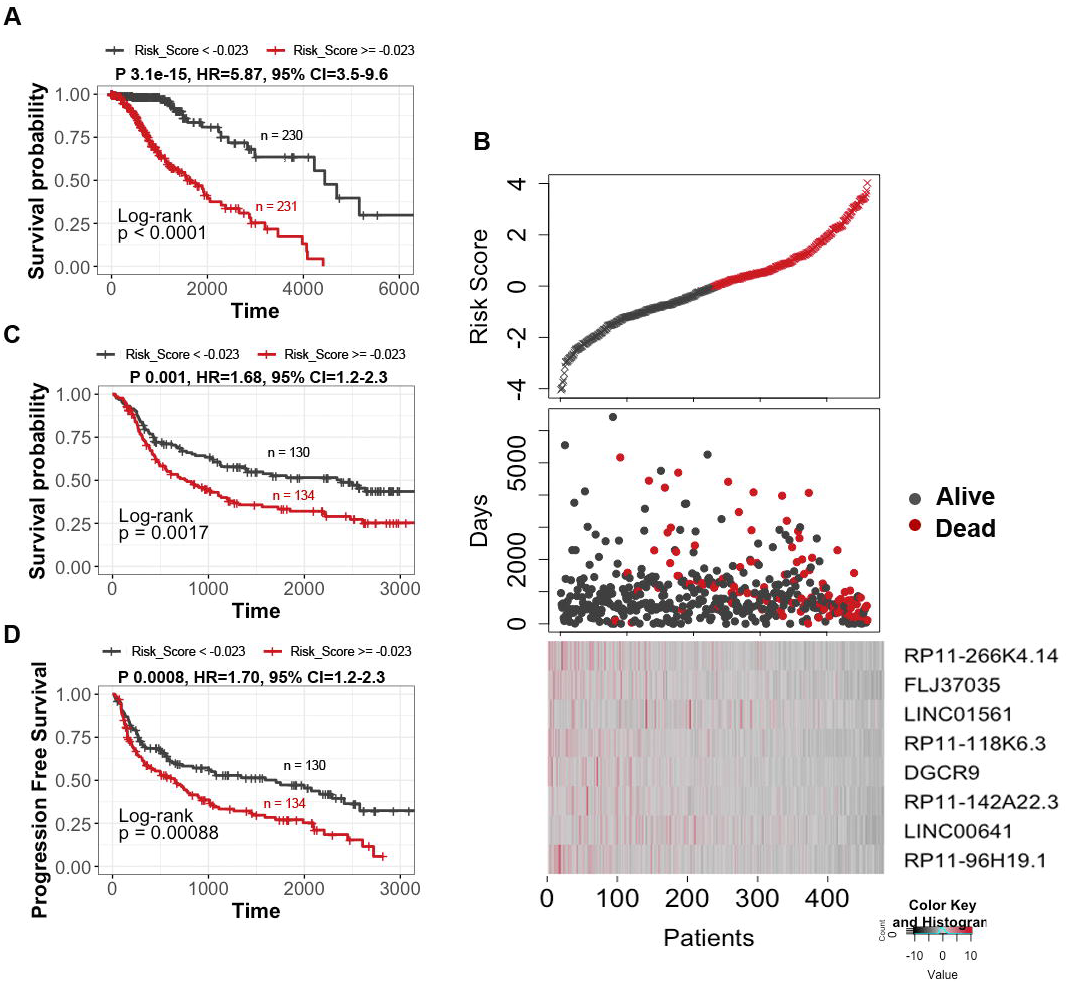
Survival analysis of the patients divided by the prognostic lncRNAs in two data sets. **A)** Patients in the entire TCGA dataset with risk score greater than median score of −0.023 show poor survival compared with patients with risk score less than median risk score. **B) Upper panel:** Plot showing patients sorted based on UVA8 risk score with black representing patient with risk score below median and red showing those with risk score above median. **Middle panel:** Number of days of survival indicated on Y-axis of patients sorted on the X-axis based on the risk scores in the top panel and alive/dead status indicated by color. **Bottom panel:** z-score transformed expression value of lncRNAs in UVA8 show higher expression in patients with low risk score. **C)** Kaplan Meier plot of overall survival of patients in CGGA dataset with risk score greater than (red) or less than (black) median risk score of TCGA dataset. **D)** Kaplan-Meier plot for progression free survival in CGGA dataset showed poor survival for patients with high-risk score. Rest as in C.

Since, 32% of patients in CGGA are in grade IV, the difference in overall survival could be due to over-representation of grade IV patients in high-risk group. However, even when only lower-grade gliomas (grade II and III) were separately examined we found significantly longer survival for low-risk versus high-risk patients (**Figure S4A**). UVA8 fails to cluster grade IV patients from CGGA into two distinct groups highlighting the specificity of signature for lower-grade gliomas (**Figure S4B**).

### 8-lncRNA based risk score is an independent predictor of survival

Lower grade gliomas have poorer outcomes in older patients, in tumors of higher grade and tumors with wild type IDH1 status (**Figure S1**). Interestingly, the risk score derived from UVA8 is higher in patients older than 40 years, patients in grade III vs grade II and patients harboring wild-type IDH1 gene (**Figure S5**). It was therefore important to determine whether UVA8 derived risk score is an independent predictor of survival. We divided the patients into younger (Age < 40) and older (Age >= 40) groups and found that risk-score can still stratify the patients into low-risk and high risk in both groups (**Figure 4A**). Similarly, UVA8 based risk score can still separate the patients into low and high-risk groups in grade II or grade III gliomas (**Figure 4B**). Although, IDH mutation status is a widely used prognostic and predictive biomarker, the UVA8 based risk score can also separate patients into two risk groups in patients presorted based on IDH mutation status **(Figure 4C)**. UVA8 derived risk score can also stratify patients into two risk groups among male and female patients (**Figure 4D**).

**Figure 4.**
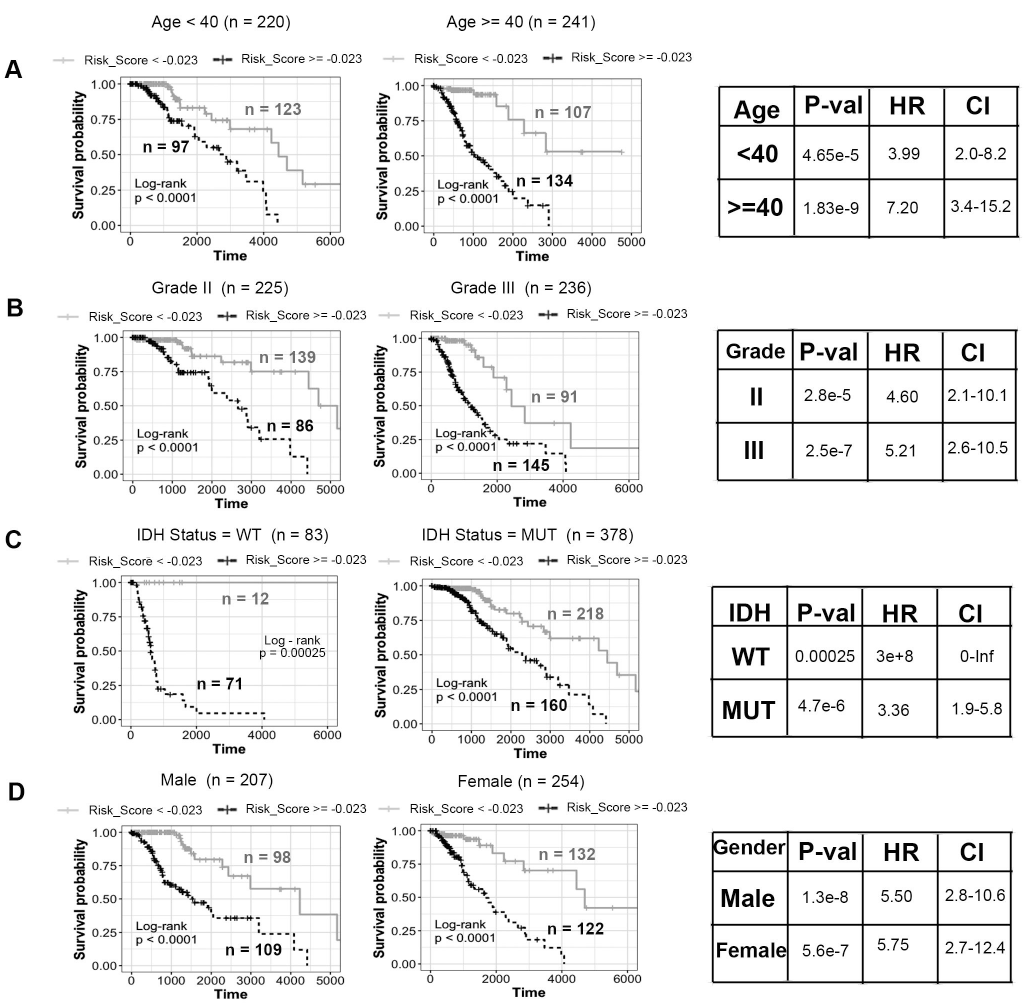
Stratification analysis by different clinical variables. Kaplan-Meier curve analysis of overall survival in high- and low-risk groups for **A)** younger (Age < 40) and older patients (Age >= 40). **B)** Grade II and Grade III patients **C)** IDH mutation status as WT and mutation (MUT) patients **D)** Male and Female patients. Black dashed line: patients with high risk score, Gray solid line: patients with low risk score. The tables on the right show log-rank p-value, hazard ratio and 95% confidence interval for each Kaplan-Meier plot.

Conversely we tested whether these standard clinically used parameters, age, gender, grade and IDH mutation status, continue to independently stratify patients even after they have been presorted into two groups by UVA8 risk score (**Figure S6**). In patients with high UVA8 risk score, age, grade and IDH mutations status can further separate the patients into two groups of better or worse outcome. In contrast, in patients with low UVA8 risk scores, none of the clinical factors could further stratify patients into two different survival groups with a pvalue<0.05 (**Figure S6**). Consistent with the previous observation (**Figure S1**), gender is ineffective in stratifying patients into two categories within patients with high- or low-risk score.

### UVA-8 is a better predictor of glioma patients’ survival

We assessed the accuracy of UVA8 in prediction of survival by comparing its time-dependent area under curve (AUC) with other clinical characteristics. For each prognostic factor (e.g. UVA8, IDH status etc.) we varied the cut-off so as to vary the false positive rate for five-year survival prediction from 0 to 1. For each cut-off the corresponding true positive rate for five-year survival was calculated (**Figure 5A**). Comparing the Area-under the curve (AUC) for these ROC curves suggested that UVA8 performs best in predicting survival of the glioma patients compared to the other criteria. This calculation was extended to predict survival of other durations (1-16 years) and the AUC plotted for each predictor (**Figure 5B**). UVA8 can predict survival better for all durations, particularly at the very early years after diagnosis when the prediction is worse for most of the predictors. Since, gender is not associated with glioma patients’ survival (**Figure S1**), the prediction of outcome was no better than random guess (AUC = 0.5) **(Figure 5A and 5B)**. We employed Cox multivariable probability hazard model to identify the impact of UVA8 and different clinicopathological characteristics in estimating hazard (**Figure 5C**). UVA8 is most significantly correlated with the survival information (p □ = □ 1.4e-07) and shows highest hazard ratio (HR □ = □ 4), indicating that the risk score performs better than any other currently used approaches for prognosis. Here, the hazard ratio of UVA8 is calculated by dichotomizing the risk score of > −0.023 (median risk score from TCGA) to 1 and < −0.023 to 0 to compare the hazard rates of high risk versus low risk patients. The hazard ratio of the 8 lncRNAs individually and combined as risk score is tabulated in **Supplementary Table S1**. The UVA8 Risk score is associated with more hazard (HR=2) than any of the individual lncRNA supporting the importance of a combinatorial signature than an individual RNA for predicting survival. The hazard ratio of UVA8 in **Supplementary Table S1** is different from that in **Figure 5C** because in the former the hazard ratio is calculated with the risk score as a continuous variable.

**Figure 5.**
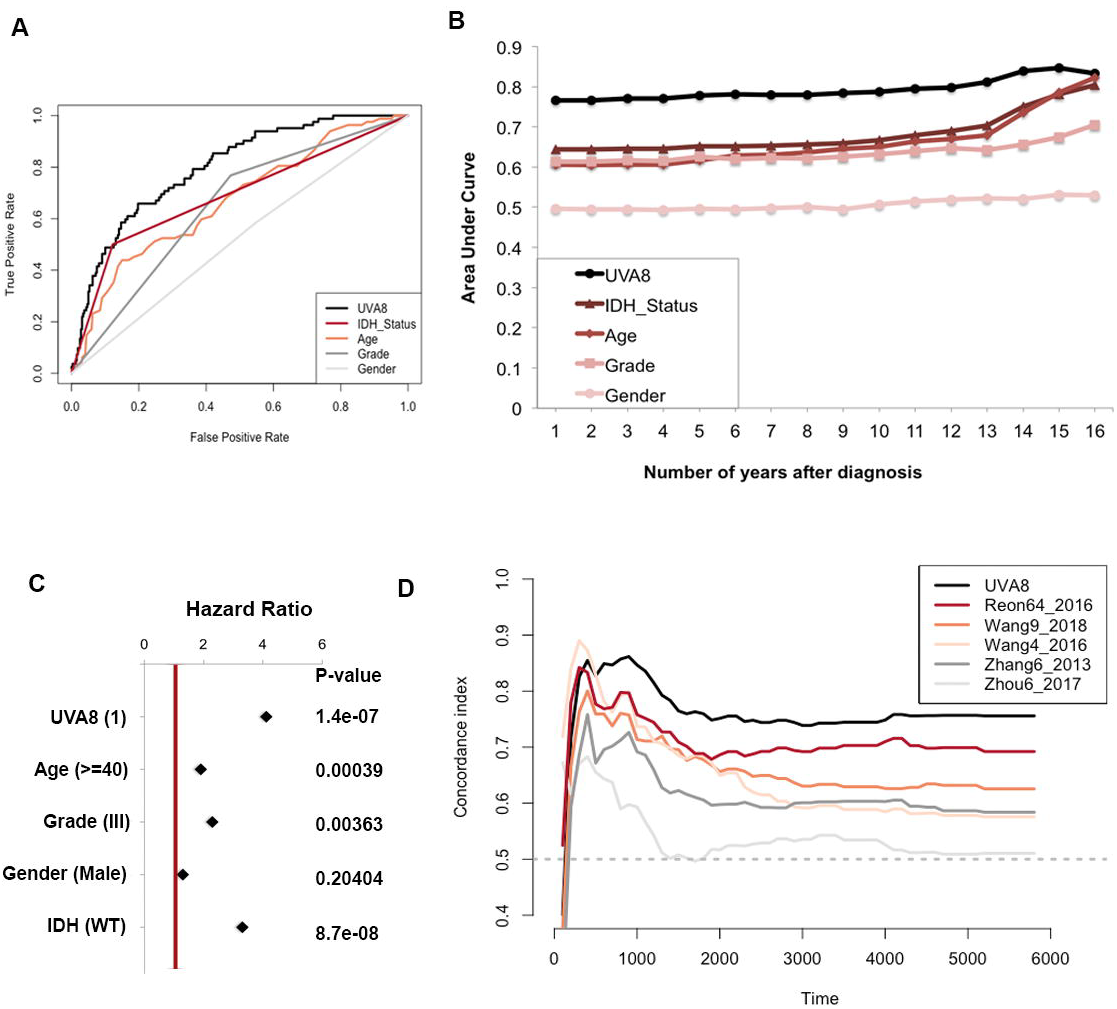
Performance evaluation of the 8-lncRNA based risk score. **A)** Receiver operating characteristic curve for 5-year survival shows UVA8 has better Area-Under-Curve compared with other predictors. **B)** Area-Under-Curve plotted for different durations of survival for 8-lncRNA based risk score, tumor grade, Age, IDH mutation status and gender of patients in TCGA cohort. **C)** Cox multivariate regression with clinical information and risk score calculated from UVA8 for survival in TCGA cohort. **D)** Concordance-index showing measure of concordance of predictor with survival of patients in TCGA.

We then sought to compare the performance of UVA8 based survival model with published lncRNA based survival models by calculating Cindex (as discussed in Methods) for TCGA dataset for each of the models. We first calculated risk score for each patient by considering the expression level of the prognostic lncRNAs in each model weighted by their estimated regression coefficients retrieved from the respective studies (**Supplementary Table S2**). The patients were ordered based on their actual survival at a given time after diagnosis and based on their risk score in each model. The concordance of the two orders is measured in pairwise comparisons of the patients to calculate a single time-dependent concordance index for the model that is being evaluated. UVA8 outperforms all existing lncRNA based survival models at different times after diagnosis (**Figure 5D).** As expected, prognostic signatures that were specific to GBMs (Zhang6_2013 and Zhou6_2017) show poor concordance index when used to predict survival of lower-grade glioma patients.

### Interferon signaling is the most enriched pathway in guilt by association with UVA8

Although many lncRNAs have been identified there has been very little functional annotation of the RNAs. We therefore applied guilt-by-association to infer functions of the lncRNAs associated with survival in UVA8. First we interrogated whether protein-coding genes most correlated with an lncRNA in TCGA glioma cohort are themselves predictive of outcome. All the lncRNAs in UVA8 are associated with a negative cox coefficient (protective). Of the 8 mRNAs most correlated positively with these 8 lncRNAs, 5 also have a negative cox-coefficient with a significant p-value. Conversely, of the 8 mRNAs most anti-correlated with these lncRNAs, 5 have a positive cox coefficient with a significant p-value (**Figure 6A**). This result is consistent with the expectation that the expression of these protective lncRNAs will be positively correlated with expression of protective mRNAs and negatively correlated with the expression of harmful mRNAs.

**Figure 6.**
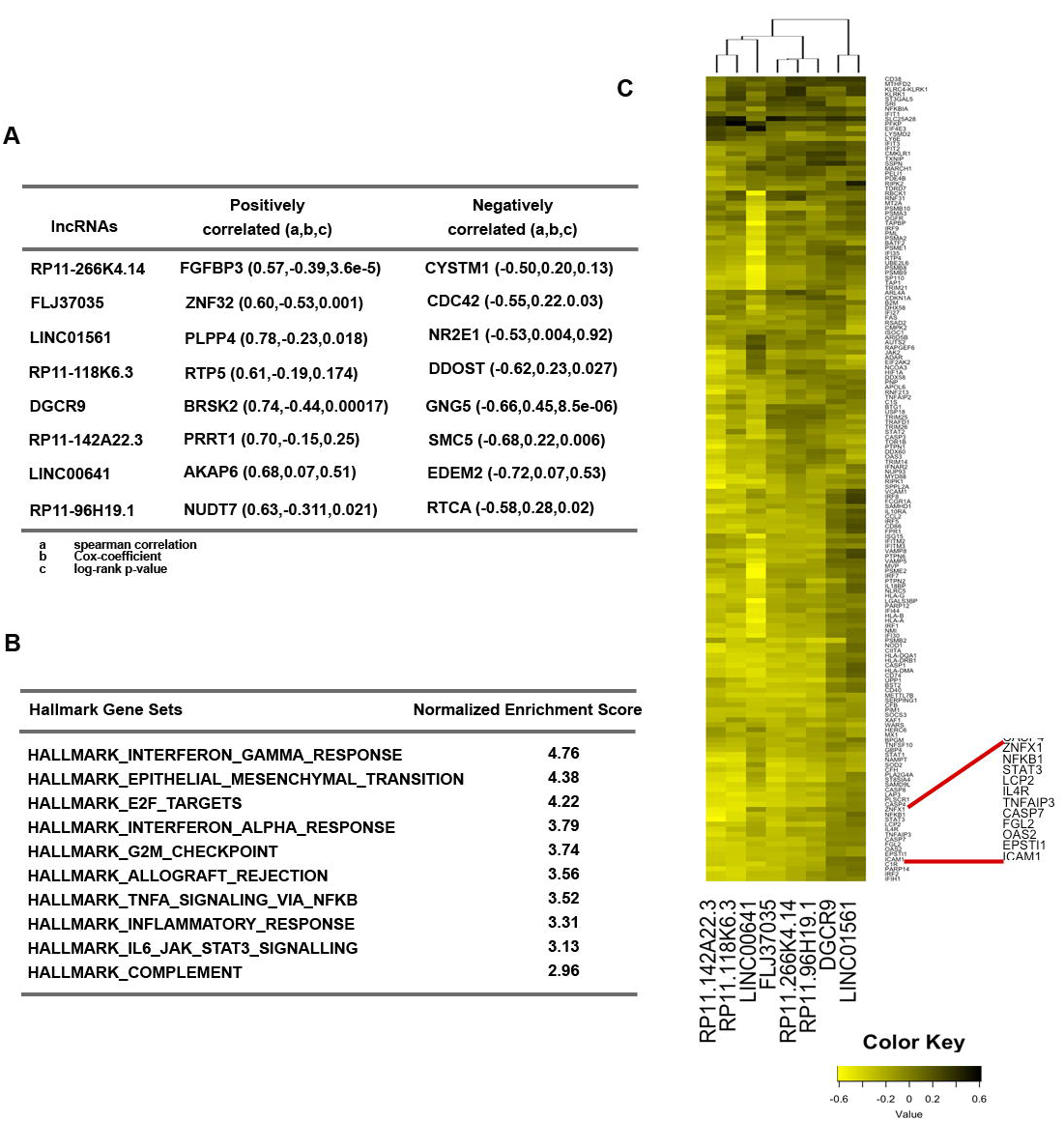
Guilt-by-association analysis of the 8lncRNAs in UVA8. A) Correlation and Cox-regression coefficient for the mRNAs that are most correlated (positive and negatively) with each of the lncRNAs in UVA8. a, b and c defined below the table. **B)** List of pathways that are most enriched in protein-coding genes that are negatively correlated with the UVA8 lncRNAs. **C)** Heatmap showing correlation of different genes in the interferon-gamma response gene set (rows) to the lncRNAs in UVA8 (columns).

GSEA analysis on protein-coding genes pre-ranked from most positively correlated to most negatively correlated to the lncRNA revealed several common pathways co-regulated with each of the 8 lncRNAs (**Figure 6B**). Interestingly, among the mRNAs that are negatively correlated with the lncRNAs, genes involved in immune and inflammatory response (IFNG, IFNA, allograft rejection, NFkB inflammatory response and JAK-STAT pathway) are highly enriched. Similarly genes involved in epithelial to mesenchymal transition and cell-cycle progressions are also most enriched. These gene-set enrichments suggest a conventional tumor suppressor phenotype associated with these 8 lncRNAs.

Many of the mRNAs are common in the IFNG, IFNA, allograft rejection, NFkB inflammatory response and JAK-STAT gene sets. The genes up-regulated in response to IFNG are mostly negatively correlated to lncRNAs in UVA8. To visualize this, the correlation coefficients were plotted for each lncRNA (columns) with individual mRNAs in the IFNG response pathway (rows) (**Figure 6C**). Out of 8, 6 lncRNAs (RP11-266K4.14, FLJ37035, RP11-118K6.3, RP11-142A22.3, LINC00641 and RP11-96H19.1) are clustered together because they are more negatively correlated with genes of interferon gamma response pathway (**Figure 6C**).

We found both NFKB and STAT3 genes as highly negatively correlated with the expression of the protective lncRNAs in UVA8. Genes involved in epithelial to mesenchymal transition and encoding cell cycle related targets of E2F transcription factors and involved in G2/M checkpoints were also negatively correlated with UVA8 expression. On the other hand, genes that are down regulated upon activation of the oncogenes KRAS are positively correlated with the expression of the protective lncRNAs of UVA8.

In order to check whether these lncRNAs can possibly act as eRNAs, we also checked the distance between lncRNAs and their correlated genes and found that these lncRNAs are correlated to several genes located in different location of genome suggesting a trans-regulation by these lncRNAs (data not shown). More experimental studies are required in future to decipher the role of these lncRNAs in regulating these genes and whether this regulation explains the effect of the lncRNAs on glioma tumor progression.

## Discussion

Gene expression profile reflects the underlying biological processes of disease. Cox regression is a widely used approach to decipher correlation between gene expression profile and patient outcome. Previous analyses on microarray data explored protein coding genes that could predict the prognosis of gliomas, particularly focusing on high grade GBMs. LncRNAs are a class of RNA which can serve as a better prognostic marker than protein coding mRNAs because they are numerous and cell-type specific [2, 3]. Additionally, since lncRNAs do not encode protein, they are the ultimate effectors, and their expression levels more accurately predict the levels of their activity. Recent studies have detected tumor-specific lncRNAs in exosomes, apoptotic bodies and microparticles highlighting another advantage of considering lncRNAs in tumors, because they are expected to appear as fluid-based markers for the diagnosis of different cancers [37–39]. Among six published lncRNA-based prognostic signatures for gliomas two are for predicting outcome in GBMs and one specifically for anaplastic gliomas. Wang et al, 2016 and Chen et al, 2017 have shown that a set of only four lncRNAs could predict survival in gliomas [23, 25]. However, the sequence of one of the lncRNAs in Chen et al., 2017, CR613436, was removed by the submitter on NCBI. Recently, the role of immune-related genes in glioma malignancies is gaining attention leading to the discovery of immune-related lncRNA-based prognostic markers for GBMs and anaplastic gliomas [40, 41]. Remarkably, there is no overlap between the prognostic lncRNAs identified in the aforementioned studies. Moreover, these studies are based on microarray data raising concerns particular to hybridization-based approaches including reliance on current knowledge of expressed genes, problems of cross-hybridization and cross-experiment comparison. Another issue is that association of lncRNAs with survival using cox-regression was sometimes carried out without controlling for any dependent variables and without penalizing for the effect of large number of variables.

In the present study, we have used an approach to screen lncRNAs from high-dimensional TCGA RNA-Seq data, which is one of the largest and the most updated data for lower-grade gliomas. After controlling for effects like age, grade, gender and IDH mutation status, we applied regularization to penalize the effect of many dependent variables and select the lncRNAs based on 100 trials. We showed the robustness of eight-lncRNA based predictor in a completely independent cohort of Chinese glioma patients. The lncRNA prognostic signature identified in the present study, UVA8, is an independent predictor of survival in TCGA glioma patients. Since UVA8 is also a better predictor than the few patient and molecular characteristics currently used for prognosis in the clinic, a simple RNA quantification will aid the physician to decide whether to adopt more aggressive therapy at the outset.

The protective lncRNAs that constitute UVA8 are negatively correlated with protein coding genes involved in interferon gamma and inflammatory response highlighting the role of immune-response genes in glioma progression. Except LINC01561, all 7 lncRNAs (RP11-266K4.14, FLJ37035, RP11-118K6.3, DGCR9, RP11-142A22.3, LINC00641 and RP11-96H19.1) are negatively correlated to most of the protein-coding genes which are up-regulated in response to interferon gamma/alpha, genes regulated by NF-kB in response to TNF, inflammatory response, and genes up-regulated by IL6 via STAT3. This suggests that an active immune reaction perhaps in response to cytokines secreted from tumor and immune cells is predictive of poor outcome in gliomas. NF-κB and JAK/STAT pathways are known to be aberrantly up-regulated in GBMs. The level of NF-κB increases as the tumors progress in astrocytic tumors [42, 43] and STAT3 is constitutively active in GBMs [44, 45]. Immune related pathways are also known to be involved in glioma tumor cell proliferation [46], survival [40], invasion [47] and chemoresistance [48]. In addition, epithelial-mesenchymal transition (associated with invasion) and active cell proliferation are suppressed if UVA8 lncRNAs are high, and this leads to better outcome, consistent with our understanding of how invasion and cell proliferation negatively impact outcome. On the other hand, genes that were positively correlated with the expression of UVA8 are enriched in genes that are down regulated by activation of the oncogene KRAS.

There are reports of the same lncRNA being predictive of outcome in the same manner in multiple tumor types. For example, DRAIC expression predicts good outcome in gliomas, melanomas, and cancers of the prostate, stomach, liver, kidney and lung [49]. In contrast, expression of LINC00152/CYTOR is predictive of poor outcome in gliomas, and cancers of the head & neck, lung, kidney, liver and pancreas (our unpublished work). Such observations are particularly exciting because they imply that the lncRNA has an important role in tumor biology that transcends tumor types, and these RNAs should be prioritized for cell- and molecular-biology studies to discern their function. It will thus be very interesting to explore whether any of the lncRNAs of UVA8 will be protective in other tumor types. Finally, future studies will address whether structural variation, copy number variations and sequence polymorphism of these lncRNAs contribute to the prognostic outcome. We are excited that UVA8 was also predictive of outcome in a completely different tumor cohort (CGGA) from a patient population that is from an entirely different geographical location with attendant differences in environment and population genotypes. It will be interesting to see if UVA8 is equally predictive of outcome in other patient populations from other parts of the world.

## Acknowledgments

We thank Dutta lab members for helpful discussions. M.K. is supported by a DOD award PC151085. The work was supported by a V foundation award D2018-002 and R01 AR067712 from NIAMS.

## Abbreviation

lncRNA: Long Non-Coding RNAs
WHO: World Health Organization
LGG: Lower Grade Gliomas
GBM: Glioblastoma Multiforme
CNS: Central Nervous System
TCGA: The Cancer Genome Atlas
CGGA: Chinese Glioma Genome Atlas
HR: Hazard Ratio
PFS: Progression Free Survival
IFNG: Interferon Gamma
Cindex: Concordance Index
AUC: Area Under Curve
ROC: Receiver Operating Characteristics

